# Effects of oxidative stress and aging on nerve, muscle, and synapse in a male-specific abdominal neuromuscular junction in *Drosophila*

**DOI:** 10.64898/2026.06.10.731480

**Authors:** Atsushi Ueda, Chun-Fang Wu

**Author notes:** Corresponding Author: Chun-Fang Wu & Atsushi Ueda Dept. of Biology, University of Iowa, Iowa City, IA 52242, USA, E-mail address.

## Abstract

Defects in *Drosophila* Cu^2+^/Zn^2+^ superoxide dismutase (encoded by the gene *Sod1*) lead to elevated oxidative stress and a drastically shortened lifespan. To contrast the effects of aging and oxidative stress on nerve conduction, synaptic transmission, and muscle excitability, we developed an easily accessible adult abdominal neuromuscular preparation, utilizing the male-specific Muscle of Lawrence (MOL) in *Drosophila*. The large size of MOL facilitated analyses of presynaptic nerve signals and postsynaptic responses that could result in sizable excitatory junctional potentials (EJPs) evoking full-blown muscle action potentials (APs) which were terminated rapidly by a characteristic afterhyperpolarization (AHP). Aged wild-type (WT) individuals (80 days or older) exhibited diminished neuromuscular transmission, mainly reflecting declines in motor axon conduction, with synaptic transmission remaining largely intact (since robust EJPs could still be evoked when nerve terminals were directly stimulated electrotonically). Additionally, muscle APs showed enhanced depolarizing peaks and weakened AHPs during current injection, suggesting weakening in repolarizing K^+^ currents. Chronologically younger *Sod1* mutants (up to 30 days) displayed similar trends of neuromuscular changes, confirming a major role of oxidative stress in aging. However, certain distinctions exist in muscle membrane properties and transmitter release machinery. A clear increase in muscle membrane resistance was seen in *Sod1* but not in aged WT. Additionally, unlike normal spontaneous release of synaptic vesicles leading to miniature EJPs (mEJPs), extremely enlarged spontaneous transmitter discharges occurred in aged WT but was never seen in *Sod1*, indicating a distinct, aging-specific alteration in transmitter release regulation. Notably, our work revealed considerable variation among individuals, ranging from transmission failure to largely intact neuromuscular functions, demonstrating the stochastic nature of functional declines due to aging and oxidative stress. Moreover, this study uncovered a well-defined common vulnerability, i.e. weakening of the Ca^2+^-activated BK current that caused drastic reduction in AHP in both aged WT and *Sod1* mutants, as confirmed by their diminishing sensitivity to the BK channel blocker paxilline, which caused striking alterations in the AHP in WT control.

## Introduction

Oxidative stress due to reactive oxygen species (ROS) has been implicated as a major contributor to systemic functional decline in aging (Harman, 1956; Harman, 1981). Free radicals, such as the superoxide anion (O_2_ ^−^), generated during normal mitochondrial metabolic processes impart oxidative stress upon cellular components (Finkel & Holbrook, 2000; Iakovou & Kourti 2022; Maldonado et al., 2023). The cytosolic enzyme Cu^2+^/Zn^2+^ Superoxide Dismutase in *Drosophila* encoded by the gene *Sod1* is an important free radical scavenger that converts superoxide into hydrogen peroxide, which is then further reduced to water by catalase (Anwar et al., 2025).

*Drosophila Sod1* mutants exhibit elevated levels of ROS and a drastically shortened lifespan and decreased locomotor ability (Phillips et al., 1989; Ruan & Wu, 2008; Şahin et al., 2017). We have previously examined the aging process of several thoracic motor circuits in wild type (WT) compared to that in *Sod1* mutant flies (Iyengar et al., 2022). The results uncovered distinct aging trajectories for different neural components in the escape reflex circuit and other motor pattern generators, demarcating age-resilient and age-vulnerable functional properties and demonstrating differential effects of aging and oxidative stress on certain circuit components. Further studies are required to elucidate the underlying changes in cellular physiology that lead to functional decline of nerve, muscle and synapse.

The neurophysiological changes associated with aging have been reported in several vertebrate preparations (Oh and Disterhoft 2020; Dobrowolny et al., 2021). However, the exact cellular mechanisms vary among preparations. Further exploration is desirable to determine the extent to which age-associated physiological changes are contributed by oxidative stress.

The *Drosophila* larval NMJ preparation has contributed much to the fundamental knowledge of cellular and molecular mechanisms in the development and function of nerve, muscle, and synapse (Budnik & Gramates 1999; Budnik & Ruiz-Cañada 2006). Our previous work in larval NMJs has contrasted effects of *Sod1* mutations with oxidant treatment (paraquat feeding, Ueda et al., 2021). However, an adult counterpart of the larval NMJ preparation would be highly desirable for proper aging studies, with which researchers can benefit from similar advantages in utilizing the available genetic and physiological tools and techniques to establish the fundamental knowledge for cellular neurophysiological changes underlying effects of aging and oxidative stress.

We have developed a novel adult abdominal neuromuscular preparation suitable for basic physiological studies of nerve, synapse, and muscle cellular properties. We employed the easily accessible Muscle of Lawrence (MOL; Lawrence & Johnston 1984; 1986) of male flies. With conspicuously large size, MOL facilitated current injection and intracellular recordings of excitatory junctional potentials (EJPs) and muscle action potentials (APs). Further, this preparation also enabled simultaneous en passant recordings of compound APs from the innervating segmental nerve bundle to correlate the pre- and post-synaptic events (Wu et al., 1978; Ganetzky & Wu 1982).

In this preparation, we were able to delineate the distinctions and similarities between the effects of oxidative stress and aging as manifested at the cellular level in *Sod1* mutant and aged WT flies. We demonstrated the similar alterations or declines in nerve conduction, synaptic transmission and muscle excitability. Nevertheless, our study also revealed distinctly altered muscle membrane property in *Sod* and unusually large discharges of spontaneous transmitter release in aged WT flies. Current injection and pharmacological treatment identified the Ca^2+^-activated BK current as a cellular mechanism particularly vulnerable to aging and oxidative stress. The weakening of BK current was indicated by a conspicuous change in the waveform of muscle APs, coupled with a loss of effect of the BK channel blocker paxilline, in both aged WT and *Sod1* mutants.

Our work established a neuromuscular preparation suitable for future functional and genetic studies on cellular and molecular properties of nerve, synapse and muscle in a variety of neuromuscular disorders. Further work could also be aimed at the male-specific function of MOL to improve our understanding of its role underlying courtship behaviors.

## Materials and Methods

### Drosophila Stocks

The *Drosophila melanogaster* stocks used were the *n108* allele of *Superoxide dismutase1* (*Sod1*, generous gift from John Phillips of the University of Guelph, originally designated as *cSOD* in Campbell et al. (1986) and Philips et al. (1989)) and a wild-type (WT) strain Canton-Special (CS). In this report, we abbreviate this *Sod1* mutation as *Sod*, which was kept as a balanced stock *Sod red*/*TM3*, *Sb Ser*. Among *Sod1* alleles previously studied (e.g., Ruan & Wu, 2008), *n108* was selected for this study because this allele has been most extensively characterized and because the homozygous flies could be more readily generated for physiological studies (Iyengar et al., 2022; Şahin et al., 2017). The *Sod* and CS stocks described here were raised at room temperature on standard medium as reported in previous behavioral (Ruan & Wu, 2008) and physiological (Ueda et al., 2021; Iyengar et al., 2022) studies.

### Dissection of adult abdomen

Adult male flies of specified ages were anesthetized on ice. A selected individual was decapitated and immobilized in the supine position with a pin inserted through the cervical opening. All legs were cut off and the fly was immersed in Ca^2+^-free HL3 saline, containing (in mM) 70 NaCl, 5 KCl, 20 MgCl_2_, 10 NaHCO_3_, 5 Trehalose, 115 Sucrose, and 5 HEPES, at pH 7.2 (Stewart et al., 1994), which minimizes muscle contraction during dissection. The abdominal caudal end was cut off to create an opening, and the abdominal body wall was cut opened along the ventral-dorsal border with micro-scissors. The internal organs were removed, and the abdominal fillet was pinned open. The Muscle of Lawrence (MOL) in dorsal A5 segment were used for recording because of their larger size although the MOLs are often covered by fat bodies and not easily visible. However, the caudal end of MOLs in A5 extends to A6 where the fat bodies could be readily removed to allow microelectrode penetration. The fat bodies in A5 were partially loosened when removing the pulsatile organ (heart), but otherwise left intact to avoid damaging the segmental nerve projection to A5. The thorax was cut off from the abdominal fillet preparation unless otherwise specified. The operation was performed in a glass-bottomed recording chamber originally designed for the larval preparation as previously described (Jan & Jan, 1976ab, Wu et al., 1978).

### Electrophysiological recordings from the adult abdominal preparations

Electrophysiological recordings were performed in HL3 saline containing different concentrations of Ca^2+^ as specified. The high Mg^2+^ concentration in HL3 saline suppressed muscle movement and greatly facilitated excitatory junctional potential (EJP) recordings. The [Ca^2+^]_o_ in recording saline varied in different experiments as specified. Paxilline was purchased from Cayman Chemical (Michigan, USA). To evoke nerve action potentials (nerve APs) and EJPs, the A5 segmental nerve was sucked up en passant with a suction electrode (5-7 µm I.D.) at a proximal site near the branch point from the fused segmental nerve trunk, which contains both left and right segmental nerves originating from the ventral ganglion (Subramanian et al & Fernandes 2017). The stimulation (0.1 ms pulses) voltage was adjusted to 2 times the threshold voltage to ensure action potential initiation. For EJP recordings, intracellular glass microelectrodes were filled with 3 M KCl and had a resistance of about 60 MΩ. EJPs and mEJPs were recorded from MOLs with a DC pre-amplifier (model M701 micro-probe system, WPI, Conn., USA, and a custom-made amplifier). Direct electrotonic stimulation of the presynaptic terminal was achieved by stimulating the segmental nerve at the entry to the dorsal muscle field with a suction pipet. To ensure the depolarizing current spread electrotonically to the nearby presynaptic terminals, long-duration stimuli (1 ms) were applied. The stimulus intensity was gradually increased from low to high voltage to obtain maximum EJP responses (Wu et al., 1978; Ganetzky & Wu 1982, 1983). Compound nerve action potentials (APs) were extracellularly recorded from the segmental nerve en passant near the dorso-ventral border with a suction pipet (Wu et al., 1978; Ganetzky & Wu 1982). Signals were picked up by a differential AC amplifier (DAM-5A, WPI, Conn., USA, and a custom-made amplifier). The compound nerve APs contain signals from all motor neurons that innervate the dorsal muscles within the abdominal segment A5, including the MOLs. Motor neuron axons innervating the ventral muscles branch off at more proximal sites and were not included in the recording.

### Electrophysiology from the larval abdominal neuromuscular preparations

Third instar larvae were dissected in Ca^2+^-free HL3 saline, and EJPs and mEJPs were recorded as previously reported (Jan & Jan, 1976A; Wu et al., 1978; Ganetzky & Wu 1982; Ueda & Wu, 2006). The EJP recordings were performed in HL3 saline containing the specified concentrations of [Ca^2+^]_o_.

### Passive and active muscle membrane properties

Muscle input resistance was determined by injecting −1 nA current to observe the resulting membrane potential hyperpolarization reached the steady state (700 ms for larvae, 140 ms for adults). The amplitude of steady-state hyperpolarization was used to compute muscle input resistance and the time for reaching ½ of the steady-state value (t ½) was also measured. MOLs, but not larval muscles, could generate action potentials, which were triggered by depolarizing current injection of the same duration. Current amplitude was increased stepwise from subthreshold to suprathreshold levels. A single microelectrode was used for current injection and voltage recording. Electrode resistance was nulled via a Wheatstone bridge circuit (Ueda & Kidokoro 1996).

### Data analysis and Statistics

To compare data between two genotypes, a Student’s t-test was carried out as specified in the figure legends. For multiple comparisons of means, we used one-way ANOVA followed by Tukey’s post hoc test. For multiple comparisons of variance, we used Bartlett’s test followed by F-tests with Bonferroni’s correction. The *p* values < 0.05 were considered to be significantly different. All statistical analyses were performed using custom-written C programs.

## Results

### A novel adult abdominal neuromuscular preparation of *Drosophila*

The development of adult abdominal musculature involves the formation of a new set of identifiable muscle fibers from fused myoblasts during metamorphosis to produce the segmentally repetitive adult pattern (Miller 1950). A set of identifiable motoneurons innervates each hemi-segment musculature. With the retention of a smaller subset of larval motoneurons, most of the adult segmental motoneurons are generated de novo during pupation from neuroblasts (Subramanian et al., 2017). The adult and larval abdominal musculature differ significantly in layout in each homologous hemi-segment (Figure 1A, segments 1-8, shaded in green).

**Figure 1.**
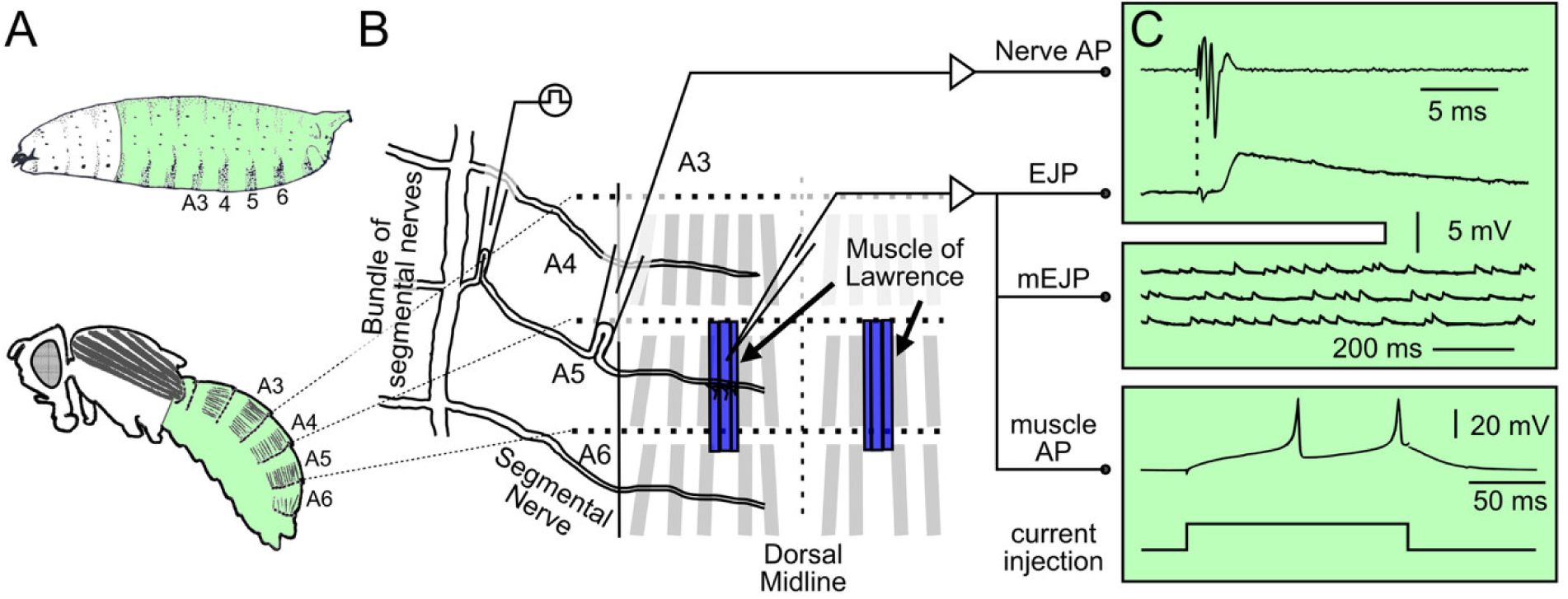
A newly developed adult abdominal neuromuscular preparation for electrophysiological studies. (A) The homologous abdominal segments between larvae and adult flies are highlighted in green. (B) Adult abdominal dorsal musculature in segments A4, A5, and A6 with the recording configuration, including a suction pipet for stimulation and another for en passant nerve action potential (AP) recordings, and an intracellular microelectrode for collecting postsynaptic excitatory junctional potentials (EJPs) and spontaneous miniature EJPs (mEJPs). The A5 segmental nerve was stimulated at a proximal site near the branch point from the fused nerve cord. This nerve cord contains both the left and right segmental nerves and originates from the ventral ganglion (Subramanian et al., 2017). In adult segment A5, the male-specific “Muscles of Lawrence” (MOLs) are conspicuous for their larger size which offers easier access to electrophysiological recordings. Note the bilateral symmetry of the musculature across the dorsal midline. (C) Electrophysiological signals obtained from the adult neuromuscular preparation. Upper panel: Simultaneous recording of segmental nerve compound APs and MOL EJPs evoked by nerve stimulation. Middle panel: Intracellular mEJPs reflecting spontaneous vesicular (quantal) transmitter release (glutamate, Rivlin et al., 2004; Hebbar et al., 2006). Lower panel: Muscle APs evoked by depolarizing current injection through an intracellular electrode using a bridge circuit (see Methods). The HL3 saline containing 20 mM Mg^2+^ and 0.75 mM Ca^2+^ (1.5 mM for muscle APs). The schematic drawings in panel A are of a larva modified from Lawrence (1992) and an adult modified from Miller (1950)

Unlike the intensively studied NMJs on larval body-wall muscles, the adult abdominal NMJs have been used mostly for morphological and developmental studies but their physiological properties have not been extensively explored (Banerjee et al., 2021; Held et al. 2019). Here we describe a filleted adult dorsal abdominal neuromuscular preparation suitable for electrophysiological studies of nerve conduction, synaptic transmission and postsynaptic muscle excitability.

Figure 1B shows a schematic of the experimental arrangements for en passant suction electrode recording of compound action potentials from the segmental nerve bundle and corresponding intracellular recordings from the male-specific Muscles of Lawrence (MOLs) in segment A5. The bilateral sets of MOLs consist of 3-5 fibers on each side (Lawrence & Johnston 1984, 1986; Gailey et al., 1991; Gailey et al., 1997; Taylor & Knittel 1995; Orgogozo et al., 2007; Kimura et al., 2024) and are known for their conspicuously large size, which afford easier access for intracellular electrophysiological recordings and current injection. We were able to collect highly reproducible and stable recordings from this special subset of adult muscle fibers. Recordings from other muscle fibers of much smaller sizes were also occasionally attempted, but with limited success. Their NMJ transmission and excitability properties may be more readily examined using different technical approaches, such as focal recording and Ca²⁺ imaging (Xing & Wu, 2018ab).

### Distinctions in spontaneous mEJP and passive muscle membrane properties between adult MOLs and larval abdominal body-wall muscles

NMJ and muscle properties in the abdominal body-wall muscles of 3^rd^ instar larvae have been well-established. Since the new preparation represents an adult counterpart that center on the male-specific MOL, we set out to explore the similarities and distinctions between the two preparations from different stages of the life cycle, using the same recording and analysis techniques.

The first notable differences were in the frequency and amplitude of spontaneous miniature EJPs (mEJPs, reflecting the release of single vesicles) between MOLs and larval muscles, with mEJPs in adult MOLs occurring more frequently (approximately 3 Hz versus 1 Hz), having smaller amplitudes (1.2 mV versus 1.5 mV), and exhibiting a faster time course (Figure 2A,C,D). We also examined the muscle passive properties to provide a basis for interpretation of the observed differences in size and time course of mEJPs. Using a bridge circuit, we were able to determine the input resistance of muscle fibers and the time constant of muscle membrane by current injection through the same intracellular recording microelectrode (Ueda & Kidokoro 1996). A step of negative current of fixed amplitude (−1 nA) produced a charging and discharging curves of hyperpolarization in both larval muscles and adult MOLs (Figure 2B), demonstrating a higher muscle fiber input resistance and a shorter half-decay time for adult MOLs compared to larval muscles 6 and 7 (14 versus 8 MΩ and 10 versus 35 ms, respectively; Figure 2EF).

**Figure 2.**
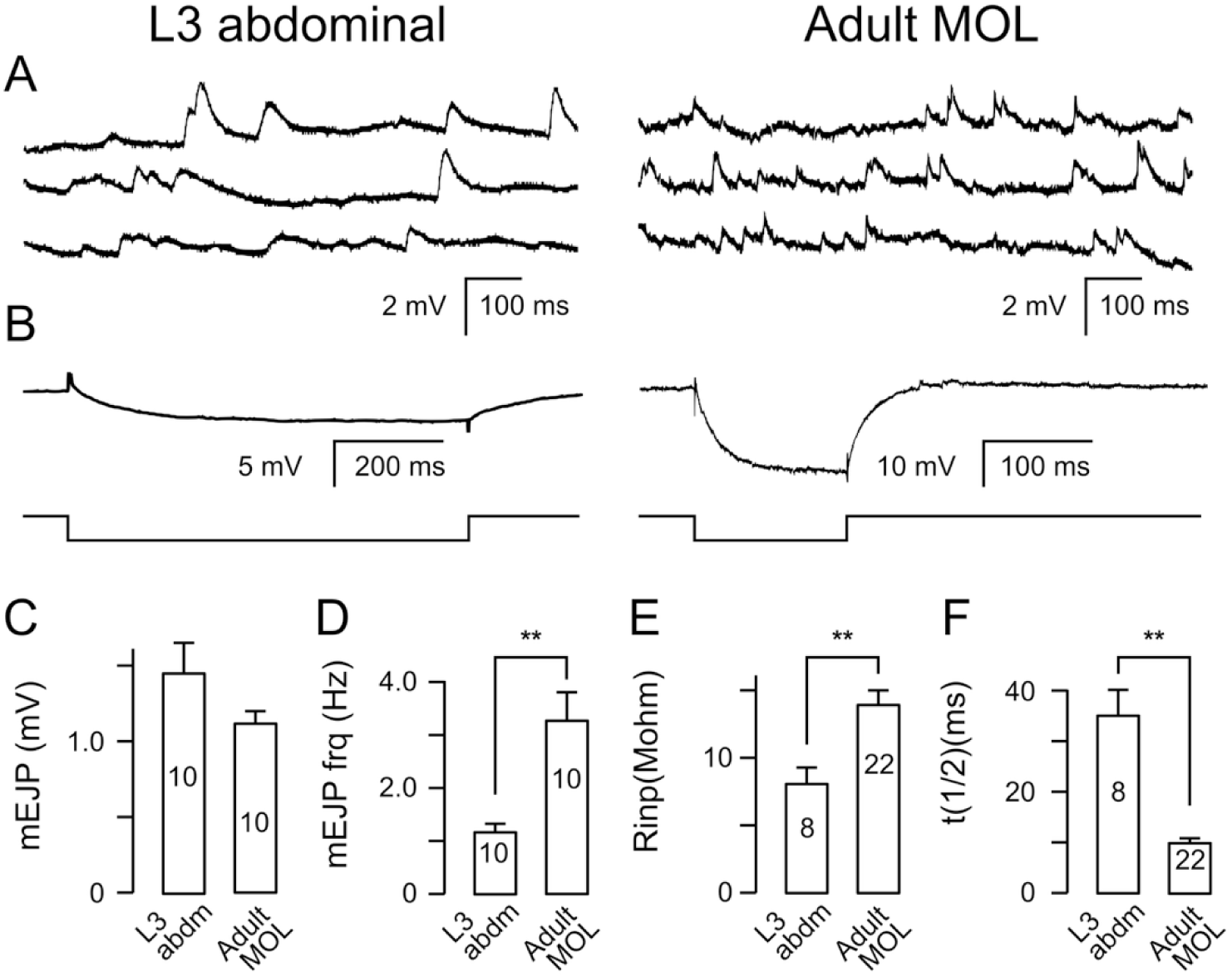
Spontaneous mEJPs and passive membrane properties measured in larval abdominal body wall muscles and adult MOLs. (A) Representative mEJPs traces from a 3rd instar larval NMJ and an adult MOL NMJ. Note that compared to larval muscles, the mEJPs in adult MOLs were smaller in size, but had a faster time course and were more frequent. (B) Electrical properties of larval muscles and adult MOLs. Representative traces of membrane hyperpolarization produced by −1 nA current injection indicate higher input resistance and a shorter membrane half-decay time in adult MOLs. Note the different time and amplitude scales. (C–F) Comparisons between larval abdominal muscles 6 and 7 (segments 3 – 5) and adult MOLs. Note the significant differences in mEJP frequency (D), muscle input resistance (Rinp; E), and muscle membrane half-decay time (t_1/2_; F). ** indicates p < 0.01 (t-test). Mean +/− SEM with sample size indicated.

The higher frequency of mEJPs in adult MOLs implies more frequent spontaneous vesicle fusion events in presynaptic nerve terminals over the muscle fiber (Figure 2D). This result could be accounted for by either more synaptic boutons innervating each fiber or more frequent spontaneous events of vesicle fusion per bouton in adult MOLs. The faster time course of mEJPs is consistent with the shorter half-decay time of the postsynaptic MOLs. However, the smaller size of mEJPs in MOLs cannot be explained by their larger input resistance, which should have resulted in larger mEJP size instead if the amount of transmitter release per vesicle and the postsynaptic receptor density and properties hold constant. Thus, smaller mEJPs in MOLs indicates either a lower vesicular load of transmitter, or different postsynaptic glutamate receptor density or conductance properties.

Assuming similar passive membrane leakage properties (specific resistance) between larval muscles and adult MOLs, the higher muscle fiber input resistance of adult MOLs is consistent with their smaller fiber size (hence total membrane area) compared to larval muscles (∼30 vs ∼80 μm width). On the other hand, the faster half-decay time of adult MOLs compared with larval body-wall muscles suggests differences in membrane properties (i.e. MOLs having a smaller value of the product of specific membrane resistance times specific membrane capacitance). Further biophysical or ultrastructural investigations may reveal the origin of the differences.

### Nerve-evoked EJPs in adult MOLs: Postsynaptic muscle regenerative potentials triggered by EJPs

*Drosophila* larvae and adults face very different environmental demands on their locomotion and movement control. Larval crawling on and digging into semi-solid food rely on coordinated peristaltic contractions of abdominal muscles (Suster & Bate 2002; Fox et al., 2006; Berrigan & Pepin 1995; Sun et al., 2022), while adult walking, climbing and flying in low-resistance air media involve the abdominal musculature with very different demands of contraction patterns and force generation. Therefore, it would be of interest to determine the distinct features of synaptic transmission and muscle response in larval and adult abdominal muscles.

We compared the EJP and muscle response waveforms at different external Ca^2+^ concentrations (Figure 3). At a lower concentration of Ca^2+^ (0.75 mM), adult MOL displayed nerve-evoked EJPs with a waveform comparable to larval EJPs, with a similar duration albeit smaller in amplitude and faster in rise time (Fig. 3Ai,ii). However, at an increased external Ca^2+^ concentration within the physiological range (1.5 mM), nerve stimulation produced in adult MOLs a response of a much greater amplitude (on average 60 vs. 5 mV), and briefer duration (less than 10 vs. tens of milliseconds). The strikingly different waveforms with a fast time course indicate activation of a regenerative muscle action potential by EJPs of increased sizes at higher Ca^2+^ concentrations (red trace in Figure 3B). It is known that action potentials in insect muscles are mediated by Ca^2+^ influx through Ca^2+^ channels (Hagiwara & Watanabe 1954; Washio 1972; Aidley 1967 for review). In contrast, larval muscles produced only a non-regenerative EJP with a prolonged duration of tens of milli-second (black trace in Figure 3B).

**Figure 3.**
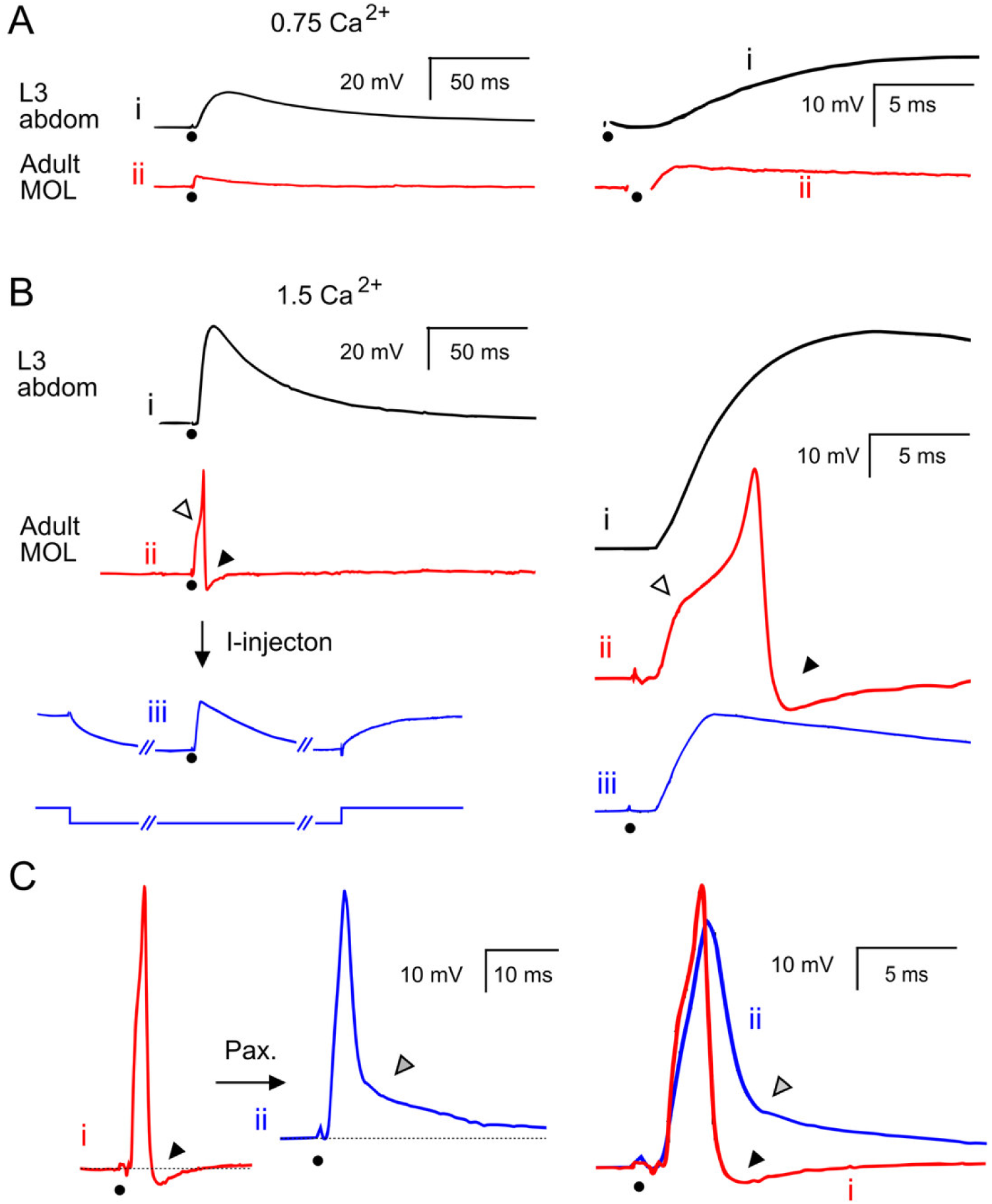
Contrasting postsynaptic responses in adult MOLs and larval abdominal body-wall muscles. Muscle EJP and AP traces shown in left panels are displayed in right panels at a faster time scale and expanded amplitude to facilitate comparisons of kinetic features. (A) At a lower external Ca²⁺ concentration (0.75 mM), adult MOLs produced a smaller EJP in response to nerve stimulation (dots) with a faster time course compared to larval muscles. The nerve stimulation artifact is blanked. (B) At an increased external Ca²⁺ concentration (1.5 mM), nerve stimulation triggered in adult MOLs a fast postsynaptic potential (PSP) of only a few milliseconds in duration, indicating activation of a fast, regenerative muscle action potential. In larval muscles, nerve stimulation evoked only a slower, non-regenerative EJP with durations of tens of milliseconds. To unmask the EJP in MOLs, the muscle membrane potential was hyperpolarized with negative current injection to prevent the nerve-evoked EJP from reaching the muscle AP threshold, revealing a graded EJP response lasting tens of milliseconds. Note that the MOL EJP triggered a muscle AP after a notch (inflection point; △) followed by a rapid repolarization due to potassium channel rectification (▴, afterhyperpolarization, AHP). Note also that the EJP during hyperpolarizing current injection lacked a notch, indicating the absence of regenerative events. (C) Adding paxilline (Pax; a BK channel blocker) to the bath widened muscle AP and eliminated the AHP (gray triangle), suggesting that the rapid repolarization and AHP involve the *slo* BK channel, which is activated by Ca²⁺ influx during the muscle AP.

In MOLs, the non-regenerative EJPs of tens of millisecond duration could be unmasked when the muscle membrane potential was hyperpolarized by negative current injection to prevent the EJP from reaching the threshold for triggering the muscle regenerative component (blue trace in Figure 3B). The right panels in Figure 3B show in an expanded time scale that the nerve-evoked EJP triggered a regenerative AP in the MOL fiber. Note a notch (red trace, open triangle) marking the inflection point near the threshold of AP initiation followed by AP peaking and a rapid repolarization, termed afterhyperpolarization (AHP), due to rectification by potassium channel activation (filled triangle). In contrast, the underlying non-regenerative EJP uncovered during the negative current injection showed a monotonic waveform, longer duration and lacked a notch and AHP (blue trace).

The time course and waveform of the AHP and its tight temporal relation with the muscle Ca^2+^ AP suggest the action of Ca^2+^-activated *slowpoke* BK channels (Elkins et al., 1986; Elkins & Ganetzky, 1988; Atkinson et al., 1998). To further investigate the possibility of a role of BK channels, we applied the blocker of Ca^2+^-activated BK channels, paxilline (1 μM), to the bath and the AHP was readily eliminated (Figure 3C, blue trace & grey triangle). The results confirm a role of BK channels triggered by Ca^2+^ influx during muscle AP in the generation of AHP.

### Effects of aging and oxidative stress on nerve conduction and synaptic transmission

Oxidative stress in *Sod* mutant flies leads to significantly shortened lifespan (Phillips et al., 1989; Ruan & Wu, 2008). The adult abdominal neuromuscular preparation enabled us to investigate neuromuscular mechanisms for aging and disease progression in an age-controlled manner. We examined nerve conduction and synaptic transmission to contrast changes in aged flies (older than 80 days, about 90 % mortality) to those in *Sod* mutant flies, as well as in younger WT control flies (< 30 days, less than 10% mortality).

We first examined the passive input resistance of the muscle fibers and the time constant of muscle membrane by step current injection (a step of −1 nA, cf. Figure 2BEF). From the charging and discharging time course and the amplitude of hyperpolarization, no difference was found in half-decay time (t1/2) for aged WT, *Sod* and WT control MOLs, indicating similar passive membrane properties. Nevertheless, MOLs in *Sod* had a significantly higher input resistance (21.6 vs. 13.4 MΩ, Table 1), consistent with the smaller sizes of *Sod* flies, hence a smaller total surface membrane area of their MOLs, which could be observed under the light microscopy images during recording and might be verified with ultrastructural techniques in further studies.

**Table 1.**
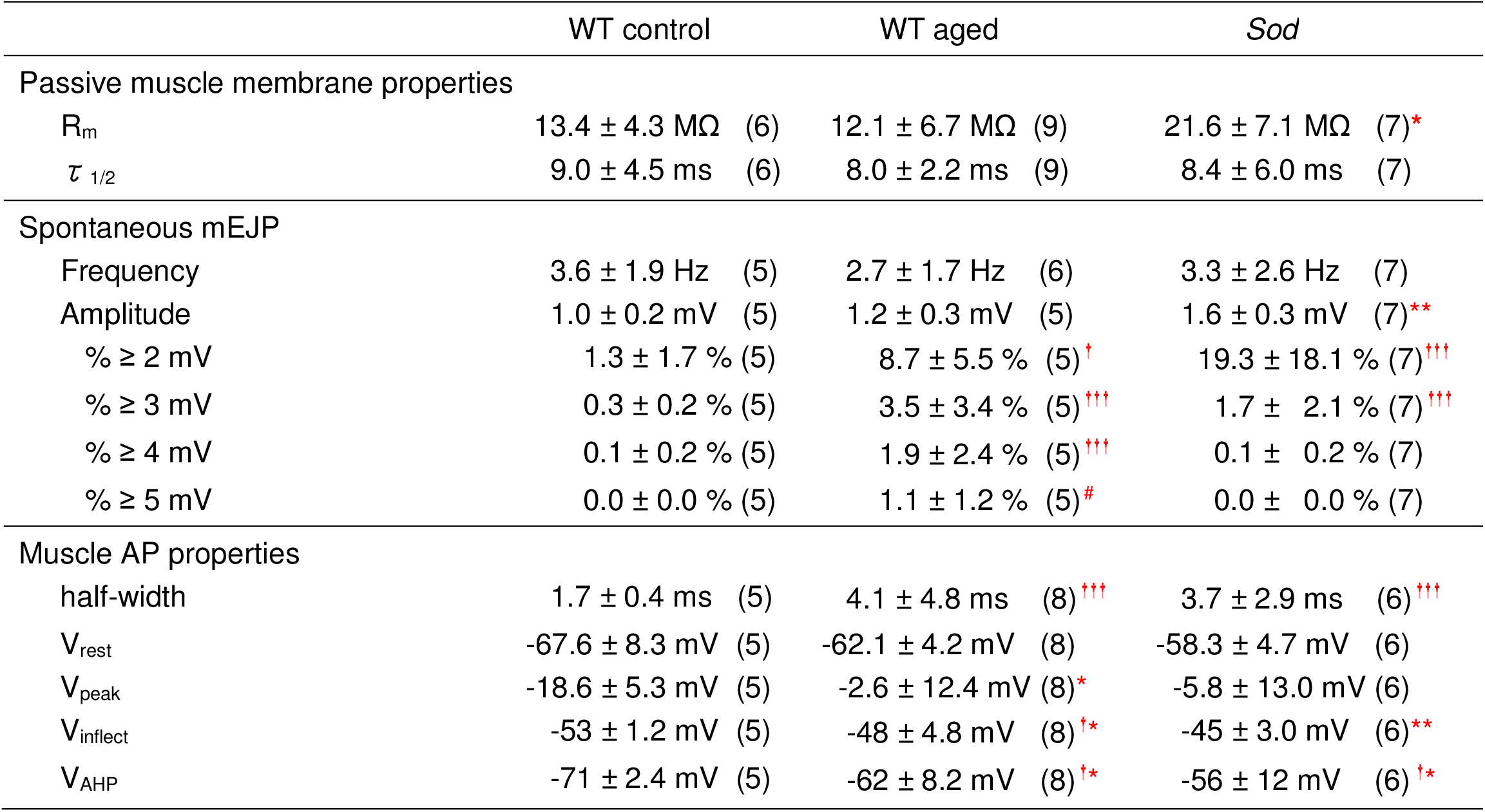
Muscle properties and miniature EJPs. WT control: 3 - 20 days old, Aged WT: 81 - 125 days old, *Sod*: 3 - 14 days old. R_m_: input resistance, τ _1/2_: charging half time, V_inflect_: Voltage at inflection point, V_AHP_: Maximum hyperpolarization of AHP, half-width: AP width at half between V_inflect_ and V_peak_. **^†^** *p*<0.05, **^†††^***p*<0.001, F-test following Bartlett’s test for variance. * *p*<0.05, ** *p*<0.01, t-test (with unequal variance when variance was different from WT control). ^#^ only aged WT showed these events (3/5 MOLs). The mEJP size distribution was non-normal, skewed toward larger size in old WT. Samples were selected for V_rest_ ≤ −55 mV. V_AHP_ was measured at the most negative potential. Mean +/− SD is shown.

**Table 2.** Functional alterations in aged WT and *Sod* mutants at MOL NMJ.

|  | WT control | WT aged | *Sod* |
| --- | --- | --- | --- |
| Nerve AP conduction | + | Weakened | Weakened |
| Nerve-evoked EJPs | + | Diminished, variable | Diminished, variable |
| Electrotonically evoked EJPs | + | Largely intact, variable | Largely intact, variable |
| Spontaneous vesicle release | ~ Normal distribution of mEJP size | Extremely skewed distribution, with some unusually large discharges | ~ Normal distribution of mEJPs with a greater mean amplitude |
| MOL AP overshoot | No overshooting (0 out of 5) | Often overshooting (4 out of 8) | Often overshooting (3 out of 6) |
| MOL AP waveform variability | Uniform | Variable | Highly variable |
| MOL AHP | Characteristic strong hyperpolarization | Weakened, variable waveform | Weakened, highly variable waveform |

We also found increases in mEJP amplitude in both aged WT and *Sod* mutant flies (Fig, 4 & Table 1). Most conspicuously, the mEJP size distribution of aged WT was highly skewed and some of the spontaneous mEJPs were abnormally large (> 500% increase). In some cases, such large mEJPs had a slower rising phase with notches, suggesting nearly synchronized multiple quantal release events (Figure 4B). However, some other giant spontaneous mEJPs lacked such notches, which and could be explained by the presence of unusually large single vesicles or certain special postsynaptic sites with greatly enhanced postsynaptic response due to altered receptor density or conductance property. The potential subcellular alterations underlying this striking phenotype may be uncovered by further ultrastructural investigations.

**Figure 4.**
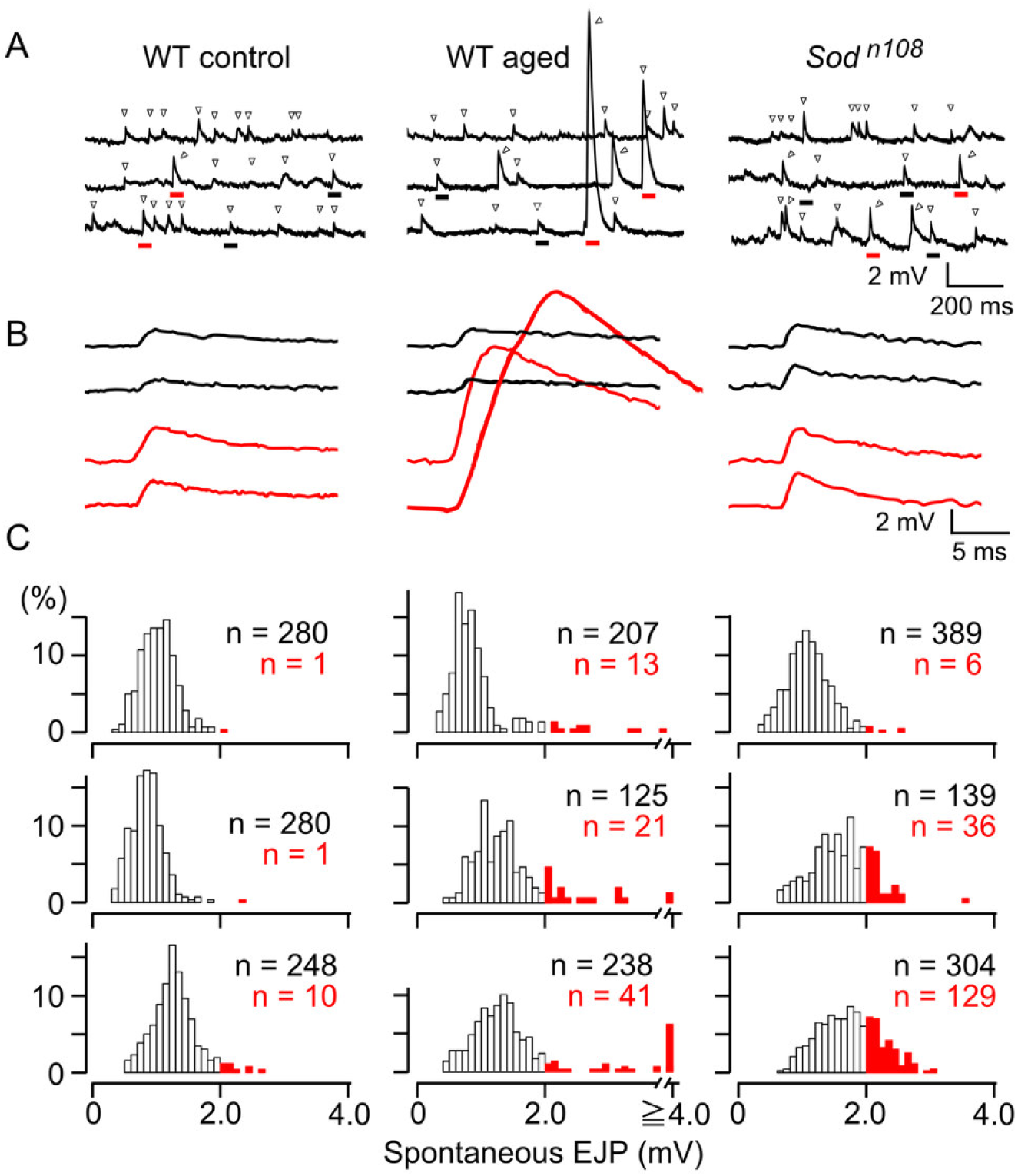
Spontaneous synaptic transmitter release in aged and oxidatively stressed flies. (A) Examples of spontaneous EJPs (▽) in younger WT, aged WT, and *Sod* mutant flies. Spontaneous EJPs in younger WT and *Sod* mutant flies were typically single-quantal miniature EJPs (mEJPs). Strikingly, aged WT flies (>80 days old) exhibited extremely large spontaneous EJPs. (B) Selected, representative spontaneous events in A (underlined) are shown enlarged at expanded scales. Events of average mEJP size are shown in black and those larger than 2 mV are shown in red (correspondingly underlined in black or red in (A). (C) Frequency distributions of the size of spontaneous EJPs shown for 3 representative NMJs of each genotype category. The red area in the distribution marks events with sizes greater than 2 mV. In aged WT, the large-amplitude fraction was prominent, and the distribution was skewed to the right. Sample sizes are indicated in black and the number of events with a size greater than 2 mV in red. Age ranges: Control (younger) WT: 2–30 days. Aged WT: 81–96 days. *Sod*: 2-30 days. HL3 saline contains 1.5 mM Ca²⁺ and 20 mM Mg²⁺.

On the other hand, the *Sod* mutant flies did not show highly skewed mEJP size distributions, with no indication of abnormally large mEJPs. Although the average size of mEJPs was larger in *Sod* flies compared to WT controls (1.6 vs. 1.0 mV, Figure 4C and Table 1), this can be accounted for by the greater input resistance of the *Sod* muscles (21.6 vs. 13.4 MΩ, Table 1). The above observation shows a distinct alteration in synaptic release machinery resulting from the normal aging process but not accelerated aging in *Sod*.

We next examined alterations in axonal function and nerve stimulation-evoked synaptic transmission (Figure 5). In both aged WT and *Sod* mutant flies, we found severely reduced amplitude in segmental-nerve compound APs, as well as greatly suppressed EJPs, some down to total transmission failure.

**Figure 5:**
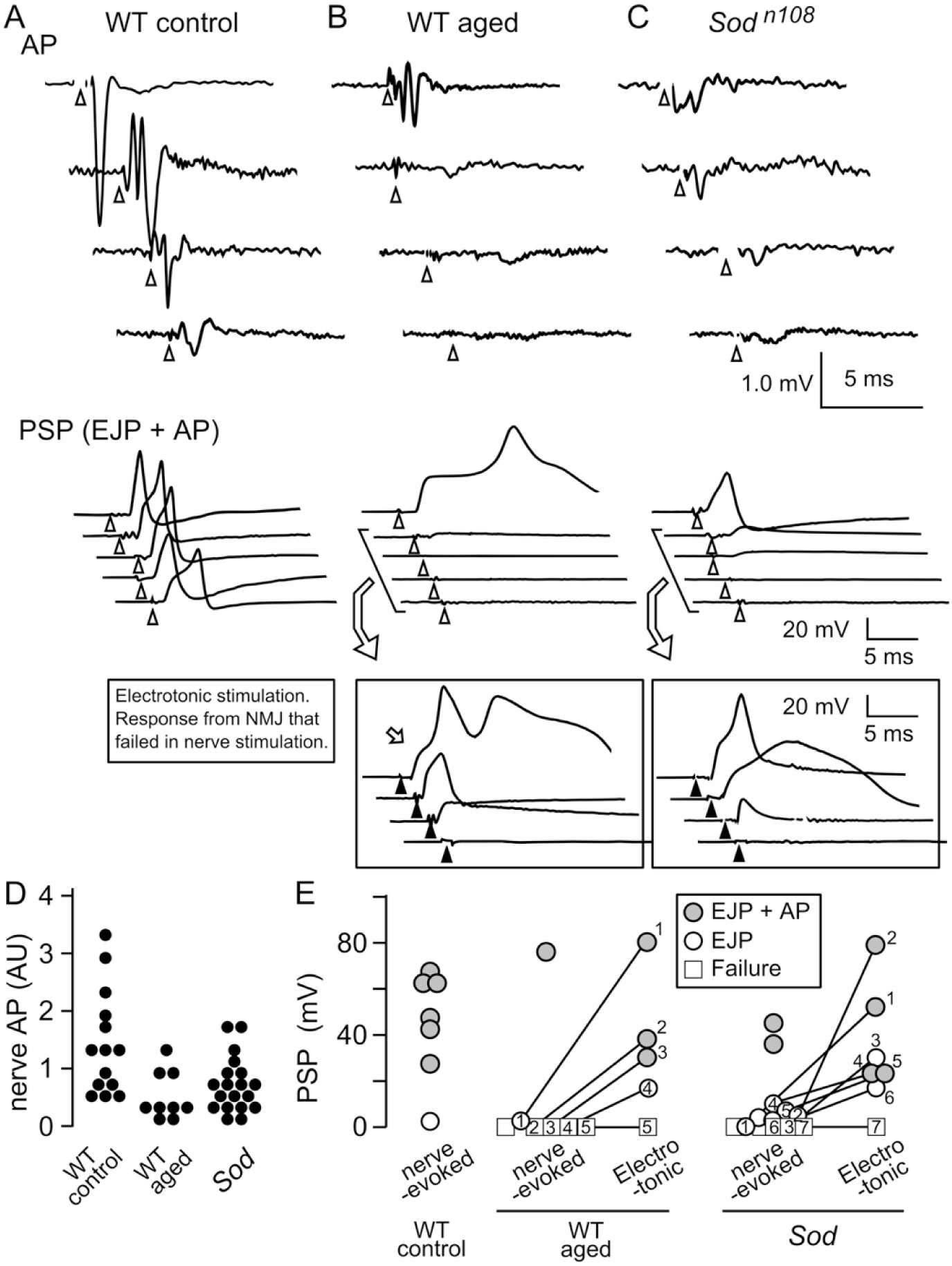
Effects of aging and oxidative stress on compound nerve APs, EJPs, and transmitter release capability in MOL NMJs. (**A-C**) Representative traces of compound nerve APs, nerve-evoked EJPs, and electrotonically induced EJPs from (**A**) WT control, (**B**) aged WT, and (**C**) *Sod* mutant flies. Note the severely diminished nerve APs (upper panels) and postsynaptic potentials (PSPs = EJP + EJP-triggered muscle AP, lower panels) in aged WT and *Sod* flies. While PSPs in WT control contained sizable EJP and muscle AP, PSPs in aged WT and *Sod* were mostly diminished and rarely produced EJP and muscle AP. For those muscle fibers that produced no response or only diminished PSPs, electrotonic stimulation (1 ms duration) was applied to directly depolarize the nerve terminal to examine its transmitter release capability. Repeated electrotonic stimulation at increasing intensities induced progressively greater responses (Boxed panels, maximum responses shown), demonstrating largely retained transmitter release capability and weakened nerve action potentials as the source for EJP failures in *Sod* and aged WT. Multiple samples from different preparations are presented, showing large variations in *Sod* and aged WT flies. Nerve and electrotonic stimulations are indicated by Δ and ▴, respectively. (**D**) Compared to younger WT controls, aged WT and *Sod* flies exhibited reduced nerve compound APs. The extracellular recorded compound AP is shown in an arbitrary unit (AU = 1 mV). (**E**) Nerve-evoked and electrotonically induced PSPs. Nerve-evoked EJPs were absent in a large proportion of aged WT and *Sod* flies (□ denotes total P□P failure; 〇 denotes smaller EJPs that failed to evoke muscle APs). Electrotonically induced EJPs were collected when nerve-evoked transmission failed to trigger muscle APs, with nerve and electrotonic stimulation data from the same NMJ numbered and connected by lines. Age ranges: Control (younger) WT: 2-30 days; aged WT: 81-96 days; *Sod*: 2-30 days. HL3 saline contained 1.5 mM Ca²⁺ and 20 mM Mg²⁺.

Measured extracellularly with a suction pipet electrode, the exact amplitude and waveform of nerve compound APs depend on the local configuration and fit between nerve and electrode, and the length of nerve loop drawn into the electrode (Wu et al., PNAS 1978; Ueda et al., 2006). Analyses of a large number of recordings indicated that the amplitude of compound nerve APs was significantly reduced in both *Sod* mutants and aged WT flies compared to the younger WT control. Further, in some *Sod* and aged WT samples, APs could not be detected at all (Figure 5B-D).

Correspondingly, most aged WT and *Sod* mutant flies showed EJP failure or only diminished EJPs well-below the threshold for triggering muscle APs (Figure 5BC, middle panels). It should be noted that the stochastic nature of aging process produced highly heterogeneous, variable phenotypes. In some rare cases of aged WT, nerve-evoked EJPs could still trigger a prolonged muscle AP with a delay before reaching the AP threshold. Similarly, in a few cases, *Sod* EJPs could trigger muscle APs with a waveform resembling the WT control. (Figure 6 and Table 1).

**Figure 6.**
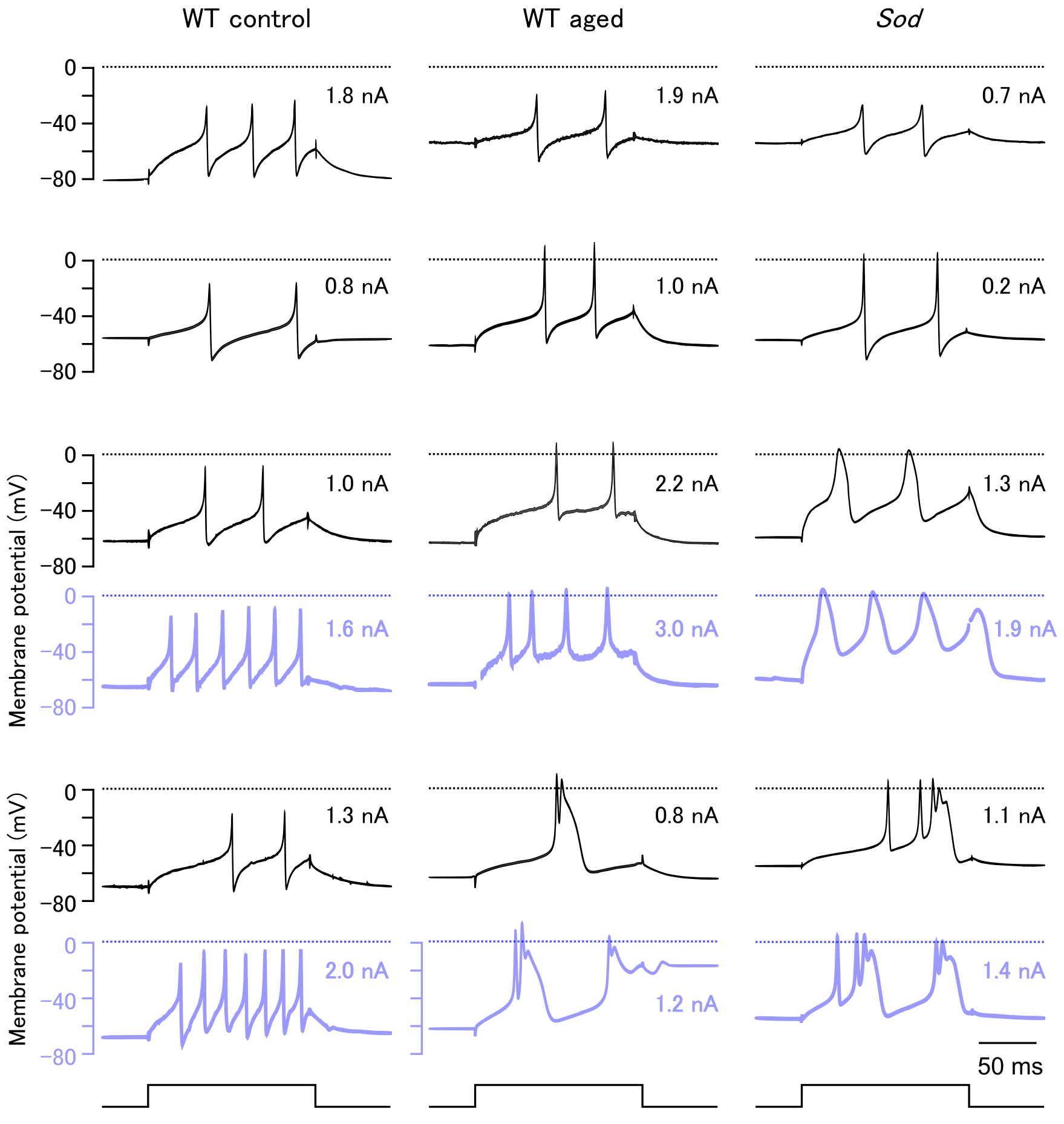
Diversity in the waveform of muscle APs elicited by depolarizing current injection in aged and oxidatively stressed flies. Spiking activities of four representative MOLs are exhibited for each category, with intensity of current injection indicated. The effect of increasing current injection intensity is shown for two cells (lower 2 panels). Note that about 50% increase lead to higher spiking rates in the same cell (blue traces). Typically, younger WT exhibited characteristic sharp APs (2-4 ms width at half maximum) without overshoot, each followed by a pronounced afterhyperpolarization (AHP). In general, aged WT produced larger APs, often overshooting but with variable AHPs. Some APs inN *Sod* muscles could resemble those in aged WT, but some others displayed much more broadened APs. The duration of step current injection is indicated at the bottom.

Considering the great variation in nerve AP phenotypes and the compounding effect on decreasing EJPs by weakened axonal action currents in aged WT and *Sod* flies, it would be desirable to examine the synaptic transmission mechanism directly in isolation of the variably weakened nerve AP. We therefore bypassed the step of motor axon AP invasion and directly depolarized the nerve terminal to induce transmitter release (Figure 5BC, boxes in lower panels). By delivering an electrotonic stimulus (longer duration of 1 ms vs. nerve stimulation of 0.1 ms) at positions closer to nerve terminals (see Methods), we were able to evoke synaptic transmitter release in most of the terminals that did not produce nerve AP-evoked EJPs in both *Sod* and aged WT flies (Figure 5E). Electrotonic stimulation with increasing intensities could lead to progressively greater postsynaptic potentials (PSPs / EJPs). Again, the maximum extent of the electrotonically induced response showed a large range of variation, from complete failure to generate postsynaptic potentials in a few terminals, indicating defective transmitter release capability, to large, prolonged PSPs, reflecting unimpaired transmitter release (Figure 5B,C,E). These large, prolonged PSPs displayed waveforms consistent with sustained muscle regenerative potentials due to weakened repolarizing forces of K^+^ channels, e.g. BK (cf. Figure 3C).

Weakened repolarization not only fails to effectively terminate muscle APs, but also enhances presynaptic terminal excitation causing prolonged transmitter release (Ganetzky and Wu, 1982; 1983) These results demonstrate that transmitter release capability could be more resilient to oxidative stress and aging effects and that weakened nerve action potential was mainly responsible for EJP failures in *Sod* and aged WT.

These observations suggest that both *Sod* and aged WT flies likely share many common defects in both nerve conduction and transmitter release mechanisms, despite the distinct giant spontaneous discharges (mEJPs) seen only in aged WT. The enormous ranges of variation in nerve conduction and synaptic transmission phenotypes displayed in both *Sod* and aged WT flies provide a striking illustration for the stochastic nature of the functional decline process in oxidative stress and aging.

### Altered muscle membrane properties of MOLs in aged WT and *Sod* mutant flies

MOLs could readily generate APs with sharp depolarizing peaks followed by a characteristic AHP, suggesting a distinct membrane excitability regulation mechanism that could be an excellent model for studying excitability property changes due to oxidative stress and aging. Upon injection of depolarizing currents as low as 1 nA, adult MOLs in WT control could fire repetitive APs when membrane depolarized above approximately −50 mV. The characteristic sharp MOL APs have a duration of approximately 2 ms (half-peak width), peaking around at −10 ∼ −20 mV (Figure 6). The membrane repolarization phase was rapid and deep, characterized by a distinct AHP of tens of millisecond duration. When a larger depolarizing current was injected, additional action potentials at a higher frequency were generated, accompanied by AHPs (Figure 6, blue traces).

Such properties of MOL APs are altered to varying degrees in aged WT flies, with different individuals displaying a range of AP waveforms (Figure 6, middle column). Most WT aged beyond 80 days produced APs with greater amplitudes, often overshooting beyond the 0 mV (dotted lines in Figure 6). AHP tended to be shallower than those in younger WT control flies (Figure 6, upper panels). These alterations suggest weakened repolarizing K^+^ channel actions.

We observed even more variable waveforms in the oxidative-stressed *Sod* MOLs (< 30 days, Figure 6, right column). Overshooting APs with long durations and different repetitive firing patterns were seen. The AHP waveform was also altered and highly variable, even more so than in aged WT flies. These alterations were more extreme in *Sod* flies but the general tendencies are largely consistent with those in aged WT, suggesting that similar changes in ion currents are induced by the *Sod* mutation and by aging in WT flies.

It is known that regenerative potentials in larval body wall muscles (Singh & Wu 1990; Gielow et al 1995; Gu & Singh 1997) and action potentials in indirect flight muscles (DLMs, Salkoff & Wyman 1980) are supported by Ca^2+^ influx through *DmCa1D* channels, whereas membrane repolarization involves actions of *Shaker* IA, *Shab* IK as well as Ca^2+^-dependent *slo* BK (Salkoff & Wyman 1980; Elkins et al., 1986; Elkins & Ganetzky 1988; Singh & Wu 1989; Haugland & Wu 1990; Wu & Haugland 1985; Ganetzky & Wu 1983; Singh & Singh 1999). Changes in these ionic currents may be responsible for the observed AP alterations in *Sod* and aged WT flies.

Both the shallower undershoot voltage for AHPs and longer duration of APs (1/2 width) in aged WT and *Sod* flies indicate weakened repolarizing forces. The Ca^2+^-activated *slo* BK current is one prominent candidate for the weakened repolarizing current because the muscle AP characteristics in aged WT and *Sod* flies can be phenocopied by treating WT control with a *slo* BK channel blocker paxilline (Figure 7; cf. Figure 2C). Reducing BK by paxilline could increase the AP peak voltage and duration and weaken the AHP in WT control, making muscle AP waveforms approaching those of aged WT and *Sod* (Figure 7A). In comparison, paxilline exerted only little or much milder effects on already altered AP and AHP waveforms in aged WT and *Sod* (Figure 7BC). It should be noted that, in some aged WT and *Sod* flies with milder AP and AHP phenotypes, application of paxilline could further increase the peak depolarization and duration of muscle AP and reduce further the extent of the AHP. These results support the conclusion that aging and oxidative stress lead to reduction in BK to variable extent and thus increase variation in AP parameters among individual flies.

**Figure 7.**
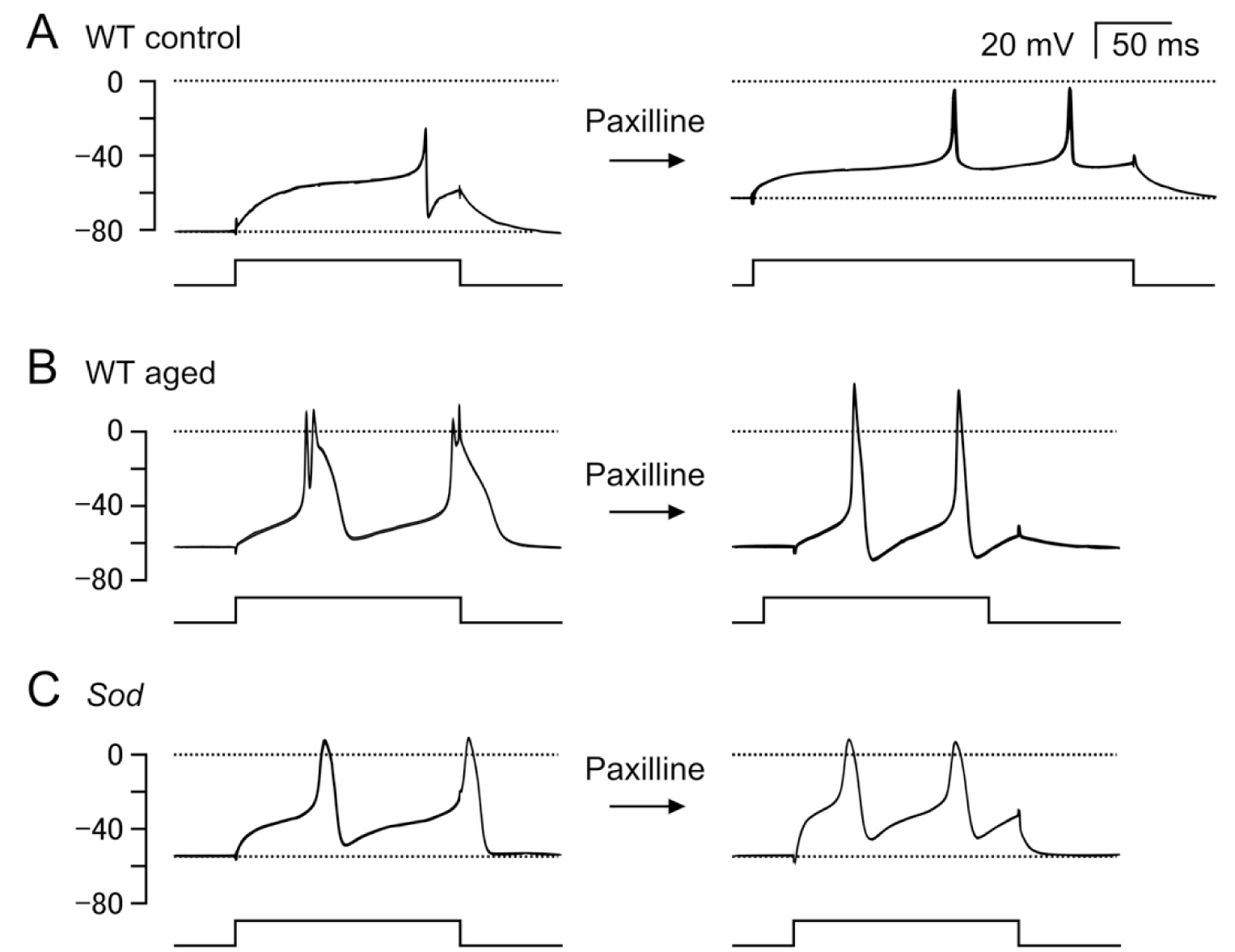
Effects of BK channel blockade on muscle APs evoked by current injection. (A) Paxilline, a specific BK channel blocker, administration to young WT reduced AHP and produced waveforms resembling phenotypes observed in some aged WT and *Sod* muscle. (B) Paxilline administration to aged WT flies modified the muscle AP waveform but did not drastically alter the slow time course of the after hyperpolarization. (C) Paxilline administration to *Sod* flies produced minimal changes to the muscle AP waveform, indicating absence of BK function that resulted in broadening of APs and replacement of the rapid AHP by slower hyperpolarizing action, presumably reflecting contributions from different K channels.

## Discussion

### A highly accessible adult neuromuscular preparation for electrophysiological analysis

We present a first description of a novel adult abdominal NMJ preparation to delineate the effects of aging and oxidative stress on basic cellular physiological properties of presynaptic axonal action potential propagation, synaptic transmission, and postsynaptic muscle cell excitability. This study was greatly facilitated by the exceptionally large size of MOLs, in which the NMJ along with the pre- and post-synaptic elements are readily accessible for electrophysiological analyses.

Our results complement the findings from several other adult NMJs that have been used to study the neurodegeneration and aging processes. These reports mostly focus on the genetic and molecular analyses of synaptic morphology and/or transmission properties, using the NMJs in a variety of muscles, including the proboscis extension muscles (Cibarial Muscle 9, or CM9; Rawson et al., 2012; Mahoney et al., 2014; Azpurua et al., 2018), abdominal ventral longitudinal muscles (Wagner 2017; Wagner et al., 2015; Hebbar et al., 2006; Banerjee et al., 2016, 2021, 2024) and dorsal longitudinal muscles (Beramendi et al., 2007), and indirect flight muscles (DLM, Martinez et al., 2007; Iyengar et al., 2022). Of these, only CM9, abdominal ventral longitudinal muscles, and DLMs have been studied electrophysiologically. Integration of discoveries of the various aspects of the NMJ in different muscles could lead to insightful further studies on age-related and oxidative stress-induced changes in physiological properties of nerve, synapse, and muscle.

The MOLs are male-specific muscles that are conspicuously large in size and thus offer several technical advantages for electrophysiological studies: (1) They are readily accessible to intracellular recording to yield stable, reproducible results. A resting membrane potential of −40 mV or deeper could be regularly obtained and maintained for tens of minutes. This allowed us to gain reliable records of evoked EJPs and spontaneous mEJPs. (2) The dorsal A5 segmental nerve innervating the MOL is long enough to allow a sufficient distance between the stimulation and recording sites, minimizing the temporal overlap between the compound nerve AP signals from stimulation artifacts, and reducing the distortion of nerve AP signals by stimulation artifacts. The long A5 nerve is also suitable for selectively applying nerve stimulation or electrotonic stimulation at appropriate sites to trigger EJPs and accompanying MOL regenerative potentials. This enabled us to analyze synaptic transmission independent of invading axonal action potential by directly depolarizing the presynaptic terminals electrotonically from the vicinity of the synapse. (3) In the MOL, a full-blown AP with a characteristic waveform can be triggered by either nerve-evoked EJP or by depolarizing current injection. This sharp AP of about 2-3 ms width was terminated by a rapid, pronounced afterhyperpolarization (AHP). By analyzing the muscle AP waveform, in conjunction with specific channel blockers, the roles of specific K^+^ currents can be established.

Male-specific MOLs are thought to be involved in courtship behavior, but their exact role remains unclear (Kimura et al., 2024; Orgogozo et al., 2007). Considering the rapid kinetics of EJPs and APs, MOLs may perform rapid movements. So far, it is known that the direct and axillary flight muscles (Ewing 1979) as well as indirect flight muscles DLMs (Ewing 1977; Salkoff & Wyman 1980) exhibit all-or-none APs. In particular, APs in DLMs are also terminated by repolarization that involves *slo* BK current (Elkins et al., 1986; Elkins & Ganetzky 1988), comparable to the AHP in MOLs (Figures 3 and 7). However, further research is needed to understand the prevalence and functional relevance of action potentials in the various muscle cells to fulfill their behavioral roles in *Drosophila*.

### Effects of aging and oxidative stress on neuron, synapse, and muscle function

Our observations demonstrated that the changes of MOLs and motor neurons induced by oxidative stress in younger *Sod* mutants were overall comparable to those associated with advanced aging in wild-type (WT) flies (80 days or older), consistent with a major role of oxidative stress in aging. Both aged WT and younger *Sod* flies exhibited significantly weakened axonal action potentials (APs) and synaptic transmission, with some individuals unable to generate detectable EJPs (Figure 5). These findings illustrate that the changes typically occurring in aged WT flies could arise in younger *Sod* flies, lending support for the idea that the aging process is accelerated in *Sod* flies (Phillips et al., 1989; Ruan & Wu, 2008), resulting in shortened lifespans of 20–40 days in different *Sod* mutant alleles compared to over 60 days in WT flies.

Notably, synaptic transmitter release capability was more resistant than axonal APs to alteration by aging and oxidative stress. This resilience was demonstrated in NMJs of aged WT and *Sod* mutant flies in which nerve stimulation ceased to evoke EJPs but transmitter release could still be induced through direct electrotonic stimulation of nerve terminals (Figure 5). At increasing intensities, electrotonic stimulation induced graded EJPs and could evoke an all-or-none muscle AP when the electrotonically induced EJP reached suprathreshold levels.

Additionally, we found that muscle APs were clearly susceptible to modification by aging and the *Sod* mutation. They exhibited enhanced peak depolarization and weakened AHP when evoked either indirectly by an EJP or directly by current injection, alterations indicating weakening of repolarizing K^+^ currents (Figure 6). Subsequent pharmacological experiments suggest that AHP is mediated by the *slo* Ca^2+^-activated BK current, which could be suppressed by paxilline, a specific BK channel blocker (Figure 7). Thus, BK channels appear to be particularly vulnerable to changes caused by the aging and by oxidative stress. Notably, APs in DLM are also repolarized by K^+^ currents including *slo* BK current (Elkins et al., 1986; Elkins & Ganetzky 1988), but whether BK in DLM is similarly altered in aging needs further investigation.

It is well established that oxidative stress can modulate ion channels (Sahoo et al., 2014; Orfali et al., 2024) and neurotransmitter release mechanisms (Giniatullin et al., 2006; Sumsion et al., 2025), not only by direct action of channel oxidation, but also via intracellular signaling pathways (Averill-Bates 2024; Koga et al., 2019).

It has been shown that *slo* BK channel could be regulated by its own oxidation (Hermann et al., 2015) and that BK channel expression can be controlled by action of cAMP-Response Element Binding protein (CREB) transcription factor (Wang et al., 2009). Since ROS signaling interacts with mitogen-activated protein kinase (MAPK) and CREB (Averill-Bates 2024; Koga et al., 2019), oxidative stress may modulate the long-term expression of *slo* BK channels. Further research is required to elucidate precisely how these potential mechanisms contribute to the changes described above in nerve, synapse and muscle caused by oxidative stress and aging.

Since additional factors other than oxidative stress have been identified in aging process in several systems (Maldonado et al., 2023; Santos et al., 2024), certain differences between the consequences of normal aging and oxidative stress could be expected. Our work documents a remarkable distinction between normal aging in WT and premature aging caused by oxidative stress in *Sod* flies. While aged WT individuals could display unusually large, multi-quantal level, spontaneous mEJP discharges, this striking phenomenon was not observed in short-lived *Sod* flies (Figure 4). This indicates that certain steps in neurotransmitter release mechanisms may undergo progressive, slow alterations to generate large spontaneous discharges in the long run toward the end of the lifespan. In future studies, it will be important to investigate whether these large spontaneous discharges are correlated with ultrastructural changes or altered protein expression patterns as described in some aging studies on other adult NMJs. For examples, changes in gross morphology (e.g. fragmentation of synaptic boutons), immunochemical staining patterns (e.g. intensity and localization), and ultrastructure and organelle appearance (e.g. active zone formation, synaptic vesicle abundance and distribution, novel multi-vesicular bodies, etc.) have been reported (Beramendi et al., 2007; Wagner et al., 2015; Banerjee et al., 2021, 2024; Mahoney et al., 2014; Rawson et al., 2012) and can be looked for in aged MOLs.

Our results also illustrated the striking stochastic nature in aging- and oxidative stress-induced functional declines. Interestingly, the phenotypes described in this report showed a surprisingly large extent of variation among individuals of the same genotype and age categories. The large variability was observed repeatedly in the physiological measurements of nerve, synapse and muscle (Figures 5 and 6). This ranged from total transmission failure to nearly intact neuromuscular transmission, due to variable degrees of axonal AP deterioration and transmitter release capability retention, and was also exhibited in the variable waveform alterations of MOL APs. Further research is needed to identify the critical steps in the underlying mechanisms responsible for the large variation of each phenotype.

Our work established a neuromuscular preparation suitable for future studies of aging and neuromuscular disorders to reveal altered properties of nerve, synapse and muscle. With the readily available genetic and pharmacological tools, it should facilitate new discoveries in cellular and molecular mechanisms. Future work could also help uncover the exact male-specific function of MOL underlying courtship behavior and beyond.

## Acknowledgments

We thank undergraduate assistants for their efforts in maintaining *Sod* and WT stocks, in particular, Tash Khan, Benjamin Jacobs, Scott Woods, Casey Inman, Lydia Luton, Madeleine Ruzicka, Esmeralda Santillan Tejeda, and Sydney Langemo. This study was supported by NIH grants R21AG047612, R01AG051513, R21NS127364, and R21NS145387.

